# miR-409-3p represses *Cited2* at the evolutionary emergence of the callosal and corticospinal projections

**DOI:** 10.1101/2021.11.03.467191

**Authors:** Nikolaus R. Wagner, Ashis Sinha, Verl Siththanandan, Angelica N. Kowalchuk, Jessica MacDonald, Suzanne Tharin

## Abstract

Callosal projection neurons are a broad population of interhemispheric projection neurons that extend an axon across the corpus callosum to connect the two cerebral hemispheres. The corticospinal tract, comprised of the axons of corticospinal projection neurons, is unique to mammals, and its full extension to the lumbar segments that control walking is, like the corpus callosum, unique to placental mammals. The emergence of these two distinct axonal tracts is thought to underpin the evolutionary expansion of complex motor and cognitive abilities. The molecular mechanisms regulating the divergence of corticospinal and callosal projection neurons are incompletely understood. Our recent work identifies a genomic cluster of microRNAs (12qF1/Mirg) unique to placental mammals. These clustered miRNAs are specifically expressed by corticospinal vs. callosal projection neurons during the molecular refinement of corticospinal vs. callosal projection neuron fate (1). One of these, miR-409-3p, can convert layer V callosal into corticospinal projection neurons, acting in part through repression of the callosal-expressed transcriptional regulator Lmo4. This conversion is partial, however, suggesting that miR-409-3p represses multiple callosal projection neuron control genes in order to specify corticospinal projection neurons. One potential additional target of miR-409-3p repression is the callosal-expressed transcriptional co-activator Cited2. *Cited2* interacts genetically with *Lmo4*, and Lmo4 can partially functionally compensate for Cited2 in thymus development(2). Further, Cited2 and Lmo4 function as opposing molecular controls over specific areal identity within superficial layer callosal projection neurons of the somatosensory and motor cortices, respectively (3). *Cited2* is highly expressed by callosal, relative to corticospinal, projection neurons from the earliest stages of neurogenesis. Cited2 is necessary for the expansion of intermediate progenitor cells (IPCs) in the subventricular zone (SVZ), and the resulting generation of superficial layer callosal projection neurons. Here we show that miR-409-3p and Cited2 interact in IPCs and in corticospinal vs. deep layer callosal projection neuron development. miR-409-3p represses the *Cited2* 3’UTR in luciferase assays. *Mirg*, which encodes miR-409-3p, and *Cited2*, are reciprocally expressed in IPCs at e15.5 by qPCR. Furthermore, miR-409-3p gain-of-function results in a phenocopy of established *Cited2* loss-of-function in IPCs. Later on, miR-409-3p and Cited2 exert opposing effects on the adoption of corticospinal vs. callosal projection neuron subtype identity. Taken together, our work suggests that miR-409-3p, and possibly other 12qF1 miRNAs, represses Cited2 in IPCs to limit their proliferation, and in developing corticospinal and deep layer callosal projection neurons to favor corticospinal fate.

## Introduction

Callosal projection neurons are a broad population of interhemispheric projection neurons that extend an axon across the corpus callosum to connect the two cerebral hemispheres. Callosal projection neurons and their associated axonal pathway in the corpus callosum are an evolutionary innovation thought to underpin the complex cognitive skills of placental mammals. The relative number of callosal projection neurons has expanded extensively throughout evolution of mammals, this increase in callosal projection neurons accounting for much of the neocortical thickness difference between macaque and human, for example (4-6). Callosal projection neurons are found throughout the layers of the neocortex. They are the predominant subtype of projection neuron in superficial layers (layers II/III), but about 20% of callosal projection neurons are found in deep layers, predominantly layer V. Deep layer callosal projection neurons are born at an earlier developmental stage than superficial layer callosal projection neurons, and are molecularly distinct. Although there are a small number of genes whose expression identify callosal projection neurons as a broad population, deep and superficial layer callosal projection neurons populations express distinct combinations of genes (7). Further, certain genes expressed by both, such as Satb2, appear to be regulated differently in deep versus superficial layer callosal projection neurons (8). Deep layer callosal projection neurons, in fact, have been posited to be an evolutionarily older subpopulation of callosal projection neurons, perhaps forming the earliest interhemispheric connections.

The deep layer callosal projection neurons of Layer V are generated concurrently with subcerebral projection neurons of layer V, predominantly corticospinal projection neurons. The corticospinal tract, comprised of the axons of corticospinal projection neurons, is unique to mammals, and its full extension to the lumbar segments that control walking is, like the corpus callosum, unique to placental mammals. The emergence of these two distinct axonal tracts is thought to underpin the evolutionary expansion of complex motor and cognitive abilities. The molecular mechanisms regulating the divergence of corticospinal and callosal projection neurons are poorly understood; however, our recent work identified a genomic cluster of microRNAs (12qF1) that are unique to placental mammals, and that are specifically expressed by corticospinal vs. callosal projection neurons during the molecular refinement of corticospinal vs. callosal fate (1). One of these, miR-409-3p, has been shown to convert layer V callosal into corticospinal projection neurons, acting in part through the repression of the callosal-expressed transcriptional regulator Lmo4. Our prior data suggest, however, that miR-409-3p represses additional callosal projection neuron control genes in order to specify corticospinal projection neurons.

One potential additional target of miR-409-3p repression is the transcriptional co-activator CBP/p300 Interacting Trans-activator 2 (*Cited2*). *Cited2* interacts genetically with *Lmo4*, and Lmo4 can partially functionally compensate for Cited2 in thymus development. Further, Cited2 and Lmo4 function as opposing molecular controls over specific areal identity within superficial layer callosal projection neurons of the somatosensory and motor cortices, respectively (3). *Cited2* is highly expressed by callosal, relative to corticospinal, projection neurons from the earliest stages of neurogenesis. Expression of *Cited2* is evolutionarily conserved between macaque and mouse (3), including in the highly expanded SVZ and superficial layers found in primates, and layer V callosal projection neurons. Cited2 is necessary for the expansion of intermediate progenitor cells (IPCs) in the SVZ, and the resulting generation of superficial layer callosal projection neurons (3). Further, forebrain-specific Cited2 conditional knockout (cKO) leads to behavioral deficits associated with human neurodevelopmental disorders (Wagner and MacDonald, 2020), highlighting the importance of Cited2 in cognitive function. Whether Cited2 is also necessary for the establishment of layer V callosal projection neurons, and whether it is repressed by members of the evolutionarily-acquired 12qF1 miRNA cluster to refine the identity of layer V corticospinal projection neurons, is currently not known.

## Results

### miR-409-3p represses the transcriptional regulator Cited2

We have previously shown that miR-409-3p represses the callosal-expressed transcriptional activator Lmo4 (1). We carried out bioinformatic analyses for additional miR-409-3p targets using the search tools miRanda (9-12), Targetscan (13-17), DIANALAB microT (18-20), and miRDB (21, 22). Each driven by a different algorithm, these predict that miR-409-3p represses a second callosal-expressed transcriptional regulator Cited2. Because Cited2 and Lmo4 are known to interact genetically during thymus development(2), and because Cited2 and Lmo4 cooperatively control callosal projection neuron areal identity, the predicted interaction of miR-409-3p with Cited2 appeared to be biologically relevant. miR-409-3p is predicted to target a single site in the Cited2 3’ untranslated region (3’ UTR) (Figure 1A). To investigate whether miR-409-3p can use this site to repress gene expression, we performed luciferase reporter gene assays in COS7 cells, as previously described (1, 23, 24). We used Cited2 reporter vectors containing either wild-type or mutated (mismatch) miR-409-3p Cited2 target sites and flanking 3’UTR sequences. We found that miR-409-3p oligonucleotides significantly repress Cited2 luciferase reporter gene expression with wild-type, but not mismatch, miR-409-3p target sequences (Figure 1B). Scrambled control miRNA oligonucleotides do not repress the Cited2 luciferase reporter gene (Figure 1B).

**Figure 1:**
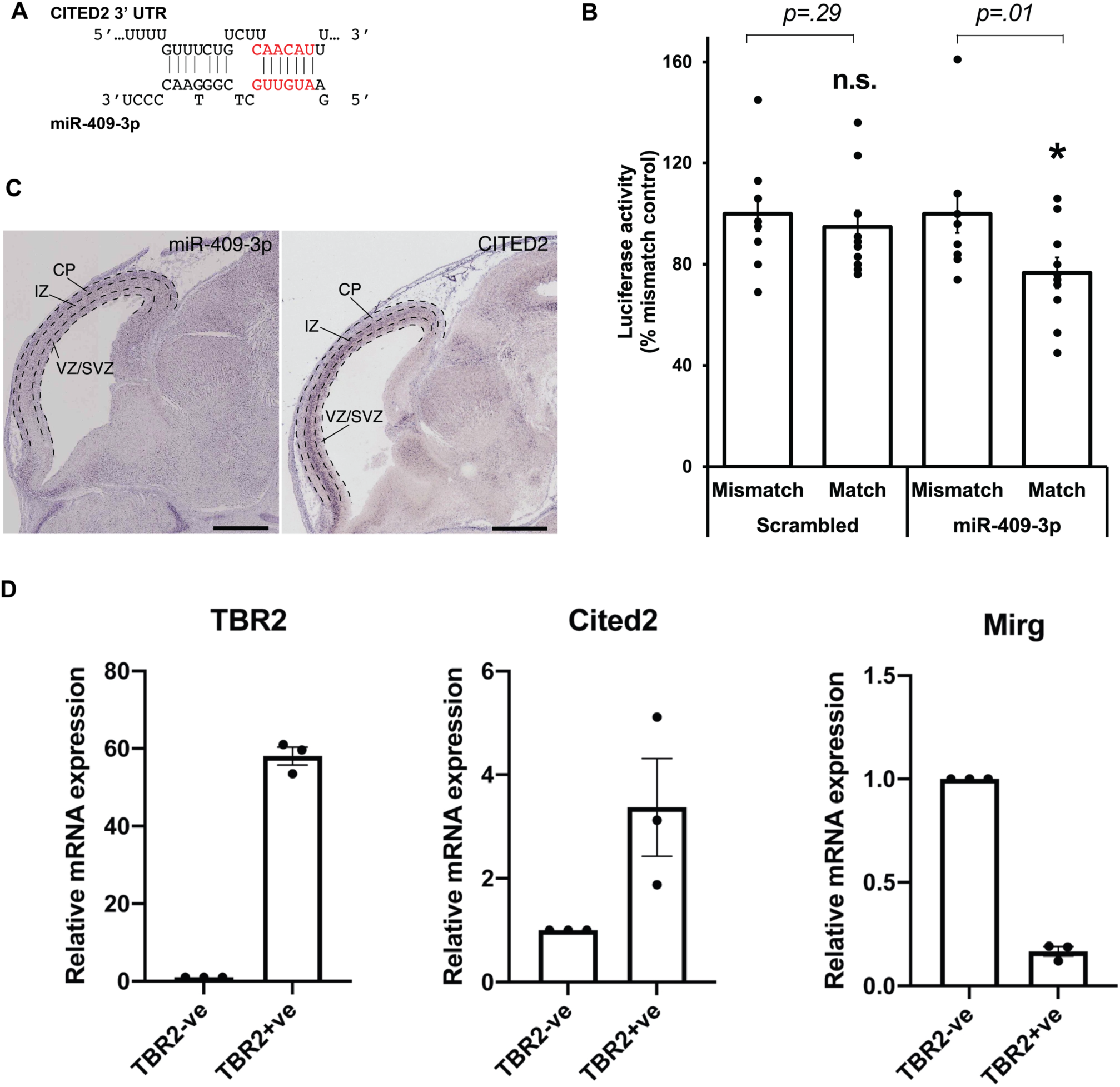
miR-409-3p represses the transcriptional regulator *Cited2*. (A) Sequence alignment demonstrating the predicted miR-409-3p target site in the Cited2 3’ UTR. Seed sequence base-pairing is in red. (B) miR-409-3p oligonucleotides repress a Cited2 3’UTR luciferase reporter gene bearing wild-type, but not mismatch, miR-409-3p target sequences. Scrambled miRNA does not repress the Cited2 reporter. Error bars represent SEM. * p<0.05 compared to mismatch control. (C) In the e14.5 dorsal telencephalon, miR-409-3p is enriched in the VZ/SVZ and CP, and not in the IZ, with minimal overlap with *Cited2*. Dashed lines indicate boundaries of the ventricular zone/sub-ventricular zone (VZ/SVZ), intermediate zone (IZ), and cortical plate (CP). Scale bars =1 mm. Raw images from the Eurexpress database (25). (D) *Mirg*, which encodes miR-409-3p, and *Cited2*, are reciprocally expressed in IPCs at e15.5 by qPCR. Error bars represent SEM. Each dot depicts an independent biological replicate.

In addition to its role in postmitotic callosal projection neuron development, Cited2 is known to be important in intermediate progenitor cells (IPCs). Specifically, Cited2 controls the number and proliferation of Tbr2+ IPCs in the neocortex at e15.5, and thereby regulates the thickness of the mature superficial neocortex (3). We determined miR-409-3p and *Cited2* expression in e15.5 Tbr2+ IPCs by purifying IPCs based on fluorescence activated cell sorting (FACS) of both Tbr2+ from Tbr2-cells from e15.5 cortices; we employed quantitative RT-PCR to measure the relative abundance of *Cited2* and *Mirg*, the mRNA encoding miR-409-3p as part of a larger, polycistronic transcript. We found that *Mirg* (encoding miR-409-3p) and *Cited2* are reciprocally expressed in these cell populations, as would be predicted if miR-409-3p were repressing *Cited2* expression in Tbr2+ cells (Figure 1D). In agreement with this finding, miR-409-3p and *Cited2* appear to be reciprocally expressed in the e14.5 developing cortex by *in situ* hybridization (Figure 1C)(25). Collectively, the data indicate that miR-409-3p can repress expression of the IPC- and callosal projection neuron-expressed transcriptional regulator Cited2, potentially thereby regulating IPC cell number and proliferation, and subtype-specific cortical projection neuron development.

### miR-409-3p gain-of-function phenocopies Cited2 loss-of-function, reducing the number of intermediate progenitor cells

We have previously shown that Cited2 is required for the expansion of IPCs in the SVZ, and that *Cited2* loss-of-function results in the generation of fewer IPCs (3). To better understand the role of miR-409-3p in IPCs, we carried out miR-409-3p overexpression gain-of-function (GOF) experiments in primary cultures of embryonic cortical neurons. Lentiviral vectors expressing miR-409-3p and GFP, and similar vectors expressing a scrambled miRNA insert and GFP were used. Cultures of e14.5 cortical cells were transfected with these vectors as described in Materials and Methods, and were examined for cell type-specific protein expression by immunofluorescence labeling on day 2 in culture. To quantify IPCs and differentiated neurons in these cultures, we labeled with antibodies to Tbr2, a marker of proliferating IPCs, to Tuj1, a marker of differentiated neurons, and to GFP, a marker of transfected neurons. Relative to scrambled control, miR-409-3p transfected cultures (GOF) display a significant decrease in Tbr2+/Tuj1-IPCs (Figure 2A,B), phenocopying the previously reported *Cited2* loss of function (3), as would be predicted if miR-409-3p were repressing Cited2 in cortical progenitors.

**Figure 2:**
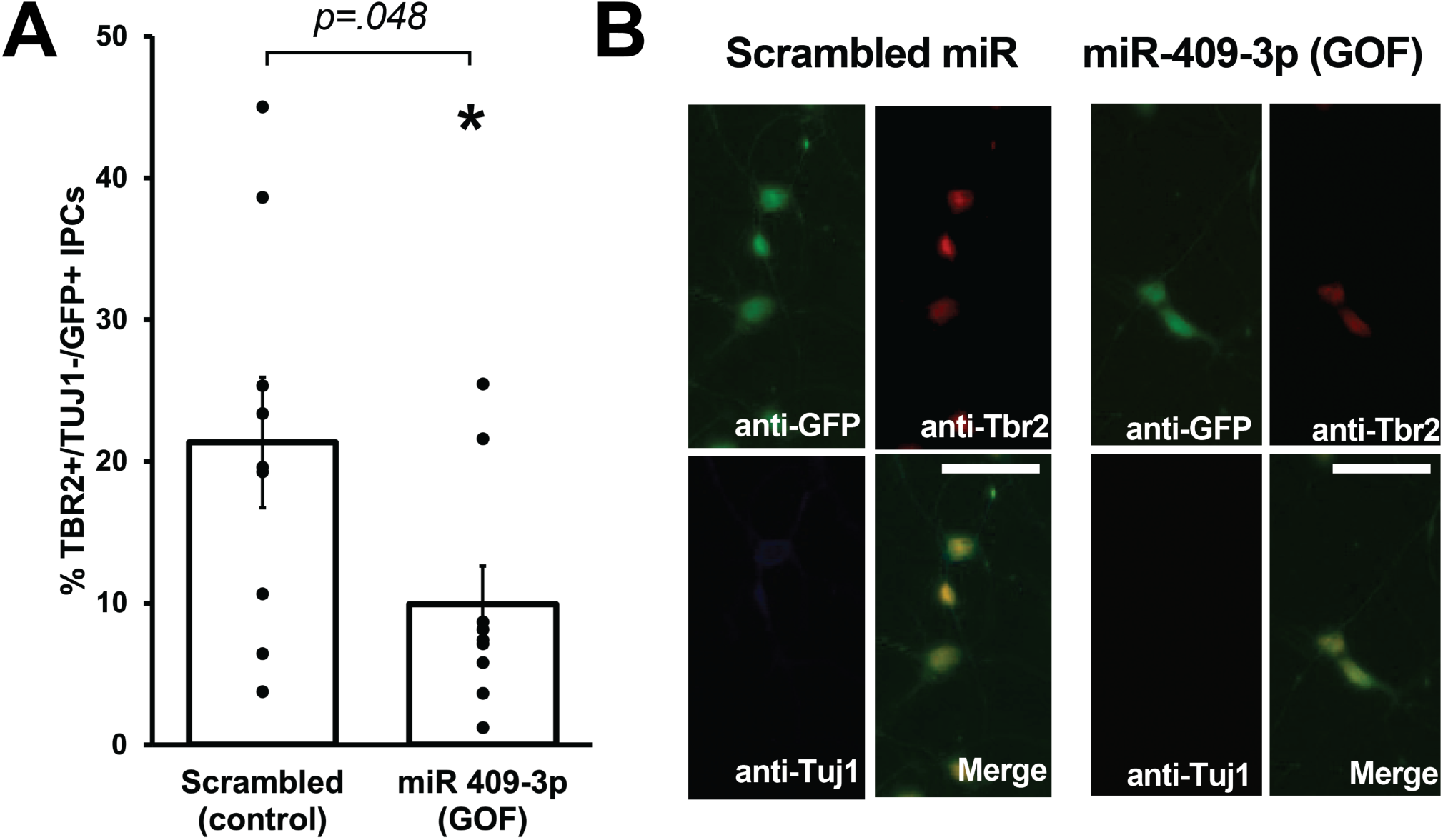
miR-409-3p represses intermediate progenitor cell proliferation in primary culture. (A) miR-409-3p overexpression gain-of-function (GOF) decreases the percent Tbr2+/Tuj1-/GFP+ IPCs compared to scrambled control in embryonic cortical cultures. (B) Representative fluorescence micrographs of embryonic cortical cultures illustrate a decrease in the percent Tbr2+/Tuj1-/GFP+ IPCs with miR-409-3p GOF. Scale bar, 50μm.

### Cited2 loss-of-function phenocopies miR-409-3p gain-of-function, promoting callosal and repressing corticospinal projection neuron identity

We have previously shown that, at e18.5, overexpression GOF of miR-409-3p (via *in utero* electroporation) promotes corticospinal projection neuron identity *in vivo* (1). In primary culture, miR-409-3p does this in part, but not wholly, via repression of Lmo4 (1). Because we suspected Cited2 was also involved in this process, we assessed the effects of Cited2 LOF on the percent corticospinal vs. callosal projection neurons *in vivo*. To do this we generated forebrain-specific *Cited2* conditional knockout (cKO) mice by crossing *Cited2* conditional floxed mice (C2f) (26) to Emx1-Cre mice (27) and homozygosing the C2f allele in the Emx1-Cre background, as previously described (3). We found that, at e18.5, *Cited2* loss-of-function increases Satb2+/Ctip2+ co-expressing cells (a subclass of developing corticospinal projection neurons), at the expense of deep-layer callosal projection neurons, specifically in developing somatosensory cortex (Figure 3A,A’). Taken together, our results suggest that miR-409-3p and Cited2 exert opposing effects on corticospinal vs. callosal projection neuron development and support our model in which miR-409-3p repressed Cited2 to favor corticospinal over callosal fate.

**Figure 3.**
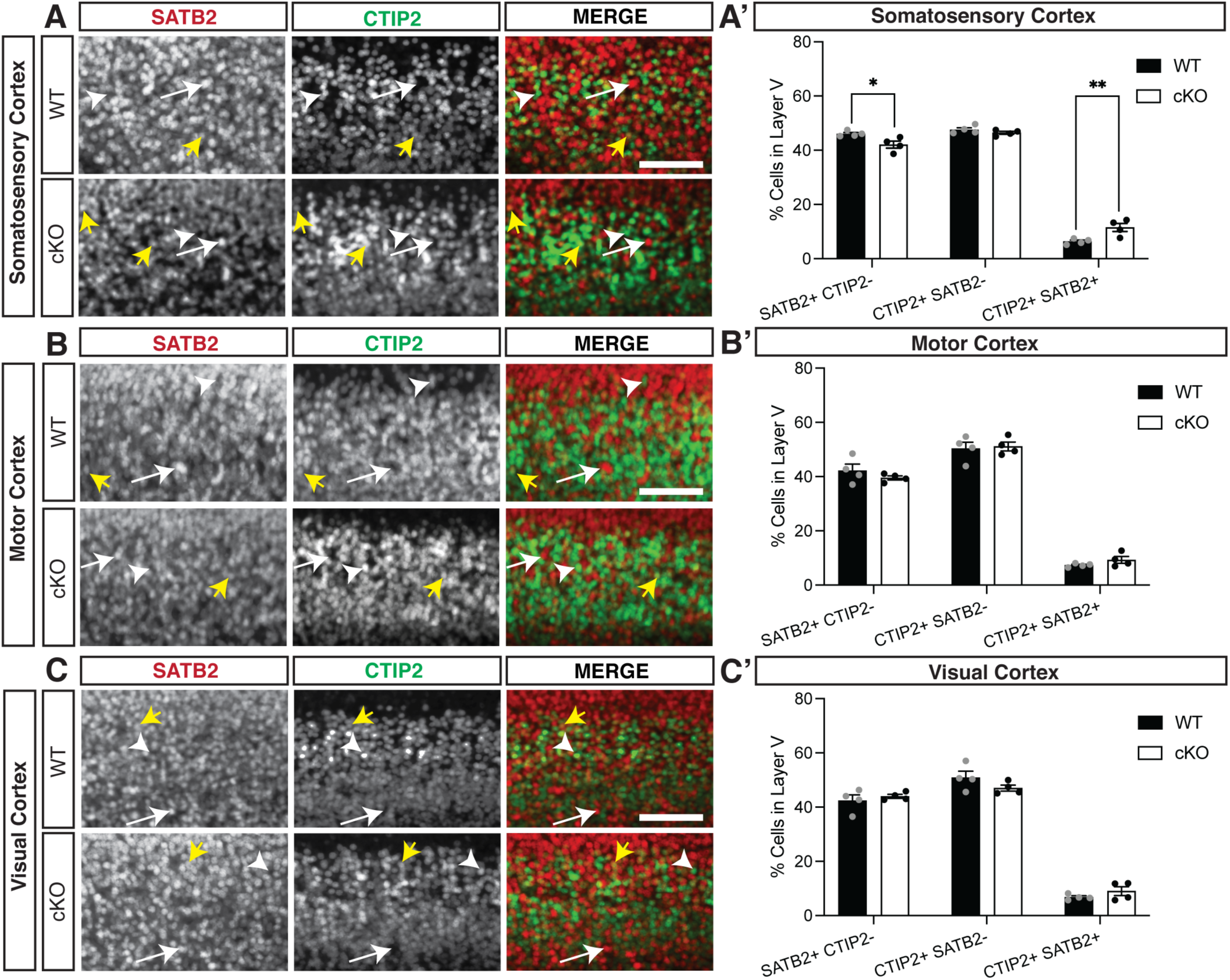
*Cited2* loss-of-function increases co-expressing SATB2 and CTIP2 cells at the expense of callosal projection neurons in Layer V of the E18.5 developing somatosensory cortex. (A-A’) Conditional loss of *Cited2* generates fewer SATB2+ CTIP2- (white arrow) neurons and more SATB2+ CTIP2+ co-expressing neurons (yellow arrows) in layer V of the somatosensory region without altering the population of CTIP2+ SATB2-neurons (white arrow head). These differences were not observed in the developing motor region (B-B’) or visual region (C-C’). n=4. *p < 0.05; **p < 0.01 (Two-way ANOVA with Šídák’s multiple comparisons test). Error bars denote SEM. Scale bars = 100 μm.

## Discussion

Corticospinal and deep layer callosal projection neurons arise from the same pool of progenitors at the same time, but they go on to adopt completely different fates in support of completely different behaviors. The key, and broadly conserved, transcriptional regulators controlling cortical projection neuron fate, while critical, wholly account for neither the narrower evolutionary conservation of the corticospinal and callosal projections, nor for their own regulation of expression. We have previously shown that a cluster of miRNAs unique to placental mammals is differentially expressed by corticospinal vs. callosal projection neurons during their development, and that one of these miRNAs, miR-409-3p, can shape corticospinal vs. callosal projection neuron fate, acting in part via the callosal-expressed transcriptional regulator Lmo4. While our findings suggested additional targets for miR-409-3p and other miRNAs in the cluster, these were up to now unknown. Here we show that miR-409-3p also targets the callosal-expressed, and Lmo4-interacting, transcriptional regulator Cited2 in cortical projection neuron progenitors. We furthermore define previously undescribed roles for miR-409-3p in controlling IPC numbers and for Cited2 in controlling callosal vs. corticospinal fate.

Because Cited2 is required for IPC expansion, the miR-409-3p/Cited2 interaction led us to investigate a possible role for miR-409-3p in IPCs. miR-409-3p and Cited2 are reciprocally expressed by e15.5 IPCs (Figure 1D), as would be expected if miR-409-3p were repressing Cited2 expression in this pool of cells. miR-409-3p GOF decreases the percent Tbr2+ IPCs in primary embryonic cortical cultures (Figure 2A,B), a phenocopy of Cited2 LOF *in vivo*(3), further supporting a functional role for miR-409-3p repressing Cited2 expression in these cells. miRNAs have previously been implicated in the production of cortical progenitors (28-46), but no role for miR-409-3p in this process was known, and none of the previously described miRNAs have been shown to target Cited2. Our findings therefore provide a link between the role for miRNAs in IPC expansion and a transcriptional regulator known to control this process.

Beyond its role in IPC expansion, Cited2 is required for the generation of superficial layer callosal projection neurons (3). However, its role in deep layer callosal vs. corticospinal projection neuron development was previously unknown. In sensory cortex, our in vivo LOF findings demonstrate that Cited2 promotes callosal projection neuron fate at the expense of corticospinal fate (Figure 3A,A’). These findings support a broad role for Cited2 in promoting callosal projection neuron development, from IPC to deep layer to superficial layer callosal projection neurons. They also further support our previously published model that miRNA repression of transcription factors that promote callosal fate in corticospinal projection neurons contributed to the evolutionary emergence of layer V projections to the corpus callosum and corticospinal tract(1).

Clustered miRNAs are known to cooperatively repress interacting genes within a pathway (47). Lmo4 and Cited2 appear to belong to such a pathway. The two genes have been shown to interact genetically during thymus development, including partial functional compensation for *Cited2* LOF by *Lmo4* in this system(2). In the developing cortex, *Cited2* and *Lmo4* appear to play region-specific roles in sculpting the areal identity of superficial layer callosal projection neurons in somatosensory and motor cortex, respectively (3). We have demonstrated that the *Mirg*/12qF1 miRNA miR-409-3p represses *Lmo4* (1) and *Cited2* (Figure 1B). Bioinformatic analyses and work by other groups suggests that multiple other miRNAs from the *Mirg*/12qF1 cluster also target *Lmo4* or *Cited2* (Figure 4).

**Figure 4.**
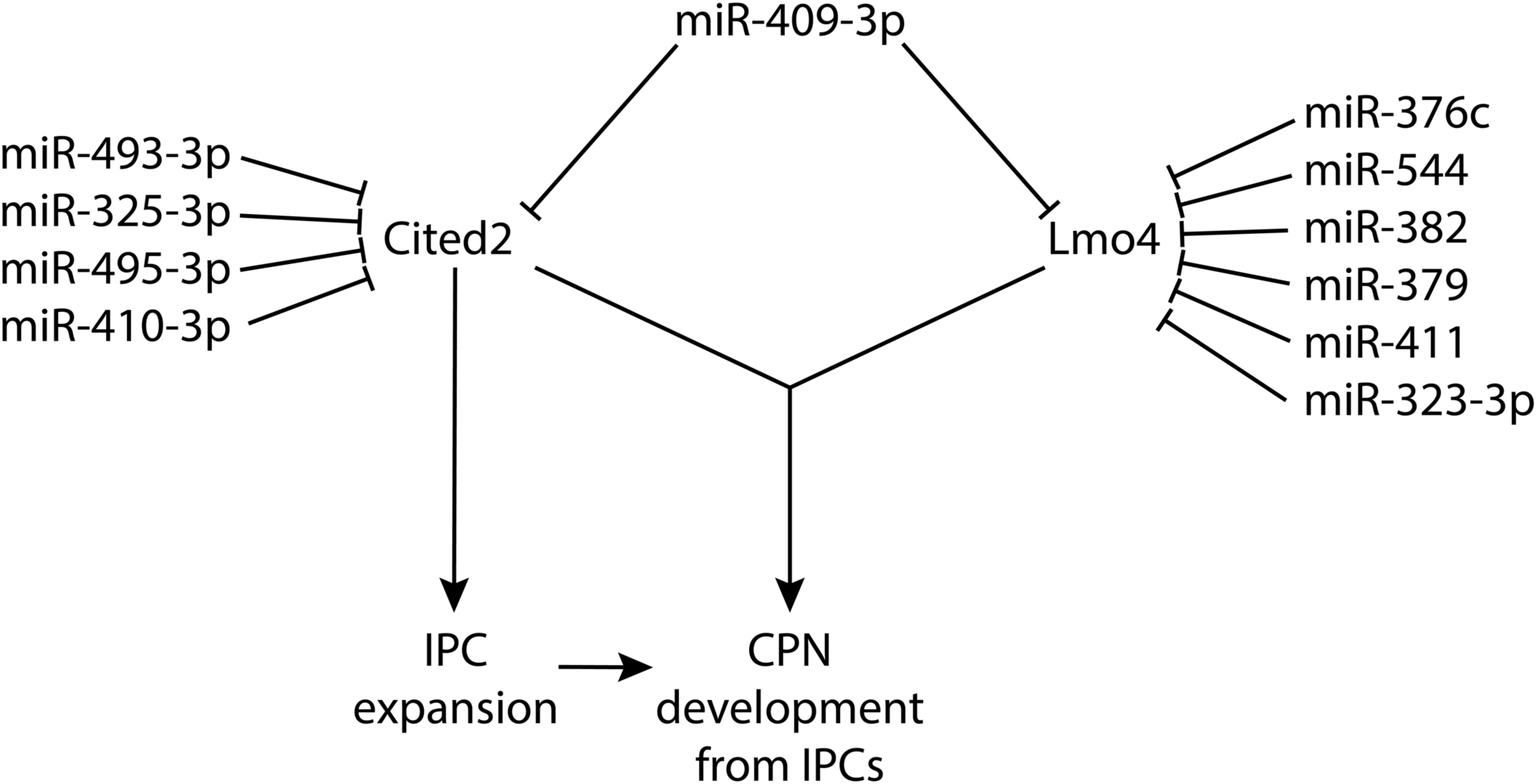
Cooperative and Multi-target repression of the Cited2-Lmo4 pathway for callosal projection neuron development by the corticospinal-expressed miRNAs of the *Mirg*/12qF1 cluster. miR-409-3p represses the interacting, callosal-expressed genes *Cited2* and *Lmo4* during cortical projection neuron development. Four corticospinal-enriched *Mirg*/12qF1 cluster miRNAs, in addition to miR-409-3p, are predicted to cooperatively repress Cited2. Six other corticospinal-enriched *Mirg*/12qF1 cluster miRNAs, in addition to miR-409-3p, are predicted to cooperatively repress Lmo4.

Clustered miRNAs are also known to cooperatively repress the same genes (47). It was previously shown that two other Mirg-encoded 12qF1 miRNAs, miR-410 and miR-495, repress Cited2 in cardiac myocytes during their development (48). Our bioinformatic analyses have identified these and two additional 12qF1 miRNAs, miR-493-3p and miR-325-3p predicted, along with miR-409-3p to cooperatively repress *Cited2*. There is thus convergent evidence supporting a role for the *Mirg*-encoded MiRNAs of the 12qF1 cluster in repressing the transcriptional regulator Cited2 during development. We propose a model whereby the clustered Mirg/12qF1 miRNAs cooperatively target the Cited2-Lmo4 callosal development pathway for repression in corticospinal projection neurons, contributing not only to refinement of dep layer projection neuron fate, but also possibly to the evolution of the callosal and corticospinal projections in placental mammals.

## Materials and Methods

### miRNA target prediction

We searched for predicted miRNA targets using the search tools miRanda (9-12), Targetscan (13), DIANALAB (18-20), and miRDB (21, 22).

### Luciferase assays

Luciferase reporter assays were performed using the Dual-Glo Luciferase Assay System (Promega), pmir-GLO based reporter constructs, and microRNA oligonucleotides (Horizon Discovery) according to the manufacturer’s instructions, as previously described (23, 24). Briefly, COS7 cells (10^4^/well) were seeded in a white 96-well plate. The following day, pmir-GLO reporter-miRNA oligo-DharmaFECT Duo (Dharmacon) transfection mixtures were prepared. The media from the 96-well plate was replaced with the transfection mixture, and the plate was incubated overnight. Firefly and renilla luciferase reporter fluorescence was read using a Tecan Infinite M1000 (Stanford High-Throughput Bioscience Center Core Facility). The ratio of firefly to renilla fluorescence was calculated for each well. Averages were compared for triplicates of each condition. Match reporter vectors contained the wild-type predicted miR-409-3p seed regions (CAACATT) with 30bp of flanking Cited2 3’ UTR on either side. Mismatch reporter vectors were identical to match reporters except that the seed sequences were replaced by GGGGGGG. Additional negative controls using empty reporter vectors and scrambled control oligos were performed. The experiments were replicated in n=5 independent cultures.

### Fluorescent Activated Cell Sorting of IPCs

Sorting of intermediate progenitor cells for qPCR analysis was based on a previously described protocol (49, 50). In brief, cortical tissue was dissected from E15.5 embryos and kept on ice in Hibernate-E (GIBCO) supplemented with 2% B27 (GIBCO) while genotyping was performed. WT and cKO cortices were then each pooled and resuspended in prewarmed digest solution in Hibernate-E minus Ca (BrainBits) containing 0.25% Trypsin (Invitrogen) and 0.01% DNase (SIGMA). Tissues were incubated at 37°C for 10 min in digest solution, washed twice in fresh Hibernate-E/B27 plus 0.01% DNase and triturated into a single cell suspension. Cells were fixed in 4% PFA plus 0.1% saponin for 30 min at 4°C, washed twice in washing buffer (0.1% saponin, 0.2% BSA in PBS), and resuspended in antibody buffer (0.1% saponin, 1% BSA in PBS) containing rabbit anti-TBR2 (Abcam Cat# ab23345, RRID:AB_778267) antibody and incubated for 1 hr at 4°C with rocking. Cells were washed twice, then incubated with Alexa Fluor goat anti-rabbit 546 antibody (1:750) for 0.5 hr at 4°C with rocking, then washed twice and resuspended in 400 μl of recovery buffer (0.5% BSA in PBS). All steps after dissociation were carried out with RNase-free reagents with solutions treated with either 1:20 RNasin (recovery buffers), 1:40 RNAsin (antibody buffers), or 1:100 RNasin (fixing solution and washing buffer). Cells were sorted at the Syracuse University Flow Core Facility with a FACS Aria II sorter (BD Biosciences) using an 85μm nozzle and FACS Diva 8.0 software. Thresholds for 546 nm sorting gates were set using secondary only controls as reference. Approximately 170,000–250,000 cells were collected for each population (TBR2+ and TBR2−) for each biological replicate. Successful sorting was validated visually using immunohistochemistry and quantified with quantitative-PCR.

### mRNA Quantitative-PCR

RNA was extracted from FAC sorted samples using Recover All™ Total Nucleic Acid Isolation kit (Ambion). RNA quantity and quality was evaluated with Agilent RNA 6000 Pico chip with a 2100 Bioanalyzer (Agilent). All RIN values were greater than 8.6. cDNA was then synthesized using qScript cDNA SuperMix (Quanta Biosciences). RT-qPCR was done using a CFX Connect Real-Time System (Bio-Rad) using primer pairs deigned to avoid genomic DNA amplification by spanning introns. Primers: *Cited2:* forward 5’ – GCT GTC CCT CTA TGT GCT G - 3’; reverse 5’ – TGG TCT GCC ATT TCC AGT C - 3’ *Mirg:* forward 5’ – TCG GCA GTA CAT ACC AGG TG – 3’; reverse 5’ – ACT GAT GGC TTC AGG TCA GG – 3’ (Sanli *et al*., 2018) *Tbr2:* forward 5’ – CAC CCA GAA TCT CCT AAC ACT G - 3’; reverse 5’ – AGC CTC GGT TGG TAT TTG TG - 3’

### Lentivirus vectors

Lentivirus vectors were modified from the pSicoR backbone (51), a gift from Tyler Jacks (Addgene plasmid # 11579). Expression of miRNA was under direction of the strong U6 promoter. miRNA inserts were either: miR-409-3p (gain of function: GAATGTTGCTCGGTGAACCCCTTTTTT), or scrambled (control, CCTAAGGTTAAGTCGCCCTCGCTCCGAGGGCGACTTAACCTTAGGTTTTT). All miRNA inserts were cloned between HpaI and XhoI sites. Expression of GFP was under direction of the CMV promoter. Lentivirus packaging was provided by System Biosciences (Palo Alto, CA). Titers of VSV-G pseudotyped viral particles were ∼10^7^ IFUS/mL.

### Cortical cultures

Embryonic cortical cultures were prepared as previously described (1). Briefly, e14.5 cortices were dissected and gently dissociated by papain digestion. A single cell suspension was prepared and plated onto poly-D-lysine (100μg/ml, Sigma) and laminin (20μg/ml, Life Technologies) coated coverslips in cortical culture medium. Cells were infected with lentivirus, and cultured on coverslips placed in 6-well plates for 2 days in growth media (50% DMEM, 50% neural basal media, supplemented with B27, BDNF, forskolin, insulin, transferrin, progesterone, putrescine, and sodium selenite).

### Immunocytochemistry of cultured cells

Cells were fixed with 4% paraformaldehyde (PFA) in PBS. Coverslips were blocked with PBS containing 0.1% Triton-X100, 2% sheep serum, and 1% BSA, and cells were then incubated with primary antibodies against cell specific markers: anti-Tbr2 (AbCam, rabbit polyclonal, dilution 1:250), and/or anti-Tuj1 (AbCam, mouse monoclonal, dilution 1:200). Following wash steps, the cells were incubated with secondary antibodies conjugated with fluorophores: anti-rat (Pierce, CY3-conjugate, dilution 1:1000), and/or anti-rabbit (Pierce, CY5-conjugate, dilution 1:1000), and anti-GFP (AbCam, goat-FITC conjugated, dilution 1:250). Cells were re-fixed in 4% PFA, and washed thoroughly in ddH2O. The coverslips were then mounted on microscope slides using Fluoroshield with DAPI (Sigma), and left overnight to dry. The following day, slides were sealed with clear nail polish, and imaged on a Zeiss AxioImager microscope. We counted transfected cells in 16 randomly selected high-powered fields, blind to experimental condition. The experiments were replicated in n=9 independent cultures. Statistical analyses were carried out in Microsoft Excel using paired two tailed t-tests with a significance threshold of p<0.05.

### Mice

All animal experimental protocols were approved by the Syracuse University Institutional Animal Care and Use Committee and adhere to NIH ARRIVE guidelines. Mice were housed at a maximum of 5 mice per cage on a 12:12 h light/dark cycle, and were given food and water *ad libitum. Cited2* conditional floxed mice (C2f) (26) were provided by Dr. Sally Dunwoodie, Victor Chang Cardiac Research Institute, University of New South Wales, Australia. They were rederived by The Jackson Laboratory (Bar Harbor, Maine, USA) using JR 664 C57BL/6 oocytes to generate live mice at an SPOF health status. Emx1-Cre mice (27) were obtained from The Jackson Laboratory (RRID:IMSR_JAX:005628). To avoid non-specific cre recombinase activity in oocytes (52), all conditional knockouts were generated by crossing C2f fl/fl females with C2f fl/+; Emx1 cre+ males, and no offspring from C2f fl; Emx1 cre+ dams were analyzed. The morning of the day of the appearance of the vaginal plug was defined as E0.5. The day of birth was designated P0. Genotypes were assessed by PCR on genomic DNA extracted with the KAPA extract kit using the following primers: *Cited2* flox/flox, flox/wt, and wt/wt were determined using – *Cited2* Forward 5’-GTC TCA GCG TCT GCT CGT TT-3’; *Cited2* Reverse 5’-CTG CTG CTG TTG GTG ATG AT −3’. Emx1 was distinguished from Emx1-Cre using – Emx1 WT Forward 5’-GAA GGG TTC CCA CCA TAT CAA CC-3’; Emx1 WT Reverse 5’-CAT AGG GAA GGG GGA CAT GAG AG-3’; Emx1-Cre Reverse 5’-TGC GAA CCT CAT CAC TCG TTG C-3’.

### Immunohistochemistry of tissue sections

Immunohistochemistry was performed as previously described (3). Briefly, E18.5 time mated brains were post-fixed overnight in 4% PFA/PBS at 4°C, then were sectioned on a VT1000S vibrating microtome (Leica Microsystems). Sections were incubated in primary antibody dilutions at 4°C overnight, and appropriate secondary antibodies were selected from the Molecular Probes Alexa series (Invitrogen, Carlsbad, CA). Antigen retrieval methods were required to expose antigens for primary antibodies; sections were incubated in 0.1M citric acid (pH=6.0) for 10 min. at 95-98°C. Primary antibodies were used as follows: rat anti-CTIP2 (Abcam Cat# ab18465, RRID:AB_2064130), and mouse anti-SATB2 (Abcam Cat# ab51502, RRID:AB_882455). Secondary antibodies from Invitrogen Alexa Series (Invitrogen). Satb2+, Ctip2+, and double positive cells were counted in Layer V over a set distance on E18.5 coronal sections matched based on subcerebral landmarks. Images were taken in the presumptive developing motor, somatosensory, and visual cortical areas based on the section’s alignment to the Atlas of the Developing Mouse Brain (53). All matching, imaging, and counting was performed by a researcher unaware of genotype. GraphPad Prism 8.0 (GraphPad Software) was used to carry out the statistical analyses. Our statistical tests consisted of Two-way ANOVA with Šídák’s multiple comparisons test to determine statistical significance between groups. Sample size and statistical test are specified in the Figure 3 legend.

